# Machine Learning Electroencephalography Biomarkers Predictive of Epworth Sleepiness Scale

**DOI:** 10.1101/2022.06.29.498173

**Authors:** Matheus Araujo, Samer Ghosn, Lu Wang, Nengah Hariadi, Samantha Wells, Saab Y Carl, Reena Mehra

## Abstract

Excessive daytime sleepiness (EDS) causes difficulty in concentrating and continuous fatigue during the day. In a clinical setting, the assessment and diagnosis of EDS relies mostly on subjective questionnaires and verbal reports, which compromises the effectiveness of available therapies. In this study, we used a computational pipeline for the automated, rapid, high-throughput and objective analysis of retrospective encephalography (EEG) data to develop objective, surrogate biomarkers for EDS, thereby defining the quantitative EEG changes in individuals with High Epworth Sleepiness Scale (ESS) (n=31), compared to a group of individuals with Low ESS (n=41) at Cleveland Clinic. Signal processing of EEG showed significantly different EEG features in the Low ESS group compared to High ESS, including power enhancement in the alpha and beta bands, and attenuation in the delta and theta bands. Moreover, machine learning algorithms trained on the binary classification of High vs Low ESS reached >80% accuracy. These results demonstrate that EEG data contain information in the form of rhythmic activity that could be leveraged for the quantitative assessment of EDS using machine learning.

## Introduction

Excessive daytime sleepiness (EDS) occurs when staying awake or alert is a progressive challenge for the individual. This condition is linked to a range of clinical factors, including metabolic and neurological diseases, but it is also associated with impairment of voluntary activities during the day or night [1]. EDS has become a significant public concern when associated with fatigue, costing more than $135 billion annually in health-related lost productivity in the United States [2]. In addition to the financial cost, the individual perception of the difficulty of concentrating and the deterioration of the brain response to audio, visual, and other stimulation motivates the search for a non-invasive biomarker that can help identify EDS to provide effective treatment. Seeking to find associations between sleepiness and its intertwined dynamics in the central nervous system (CNS), we here tested the hypothesis that EEG data contain information in the form of rhythmic activity that could be leveraged for the quantitative assessment of EDS using machine learning.

Day-time sleepiness affects the CNS and leads to changes in brain function and rhythms. Previous studies reported desynchronization between the left and right hemispheres under mental fatigue [3] and imaging data suggest altered functional connectivity between thalamus and cortex [4]. In fact, EEG has shown promising results in identifying EDS biomarkers, especially for fatigue versus alert state classification during activities such as driving using portable EEG devices [5] and for predicting driving reaction time [6]. In the clinic, subjective sleepiness symptoms can be assessed by the Epworth Sleepiness Scale (ESS), which is the state-of-practice self-report to quantify EDS or dozing propensity, and in general, it is highly correlated to standard of care sleepiness measurements such as the multiple sleep latency test [7]. However, current diagnoses methods remain essentially subjective because they rely on questionnaires and verbal reports.

In this study, we recorded resting state EEG from awake human subjects, pre-processed the EEG data using an automated artifact detection algorithm that our team has previously developed [8], followed by a statistically-guided approach for the selection of EEG features to train machine learning algorithms on the binary classification of low versus high EDS.

## Methods

### Study population

We leveraged the Cleveland Clinic Sleep Registry, a collection of multimodal physiologic data, including continuous overnight EEG monitoring housing sleep studies collected over the last decade. Data from this biophysiological repository was extracted for the purposes of this study with a focus on overnight polysomnograms or split night sleep studies. We abstracted polysomnogram data from a merged initiative of Cleveland Clinic’s distributed sleep centers to capture those with severe hypersomnolence and those without symptoms of EDS. Specifically, we identified a total of 72 patients, where n=31 with severe EDS defined by an ESS>20 and n=41 without EDS defined by ESS < 5 to serve as the analytic sample for the work presented in this manuscript. Detailed demographic information, sleep characteristics and medical history for overall patients and their subsequent division in high ESS and low ESS groups are shown in Table 1.

**Table 1.** Patient Characteristics.

|  | Overall<br>(N=72) |  | Low ESS<br>(N=41) |  | High ESS<br>(N=31) |  | p-value |
| --- | --- | --- | --- | --- | --- | --- | --- |
|  | N |  | N |  | N |  |  |
| Age (yrs) | 72 | 54.4 ± 14.5 | 41 | 55.7 ± 14.5 | 31 | 52.7 ± 14.5 | 0.39<sup>a1</sup> |
| Gender, Female | 72 | 35 (48.6) | 41 | 16 (39.0) | 31 | 19 (61.3) | 0.061<sup>c</sup> |
| Ethnicity, Hispanic | 72 | 9 (12.5) | 41 | 5 (12.2) | 31 | 4 (12.9) | 0.99<sup>d</sup> |
| Race | 72 |  | 41 |  | 31 |  | 0.097<sup>c</sup> |
| White |  | 37 (51.4) |  | 25 (61.0) |  | 12 (38.7) |  |
| Black or African American |  | 25 (34.7) |  | 10 (24.4) |  | 15 (48.4) |  |
| Other |  | 10 (13.9) |  | 6 (14.6) |  | 4 (12.9) |  |
| Sleep Procedure Type | 72 |  | 41 |  | 31 |  | 0.92<sup>c</sup> |
| PSG |  | 19 (26.4) |  | 11 (26.8) |  | 8 (25.8) |  |

Table 1 - Patient Characteristics
|  | Overall<br>(N=72) |  | Low ESS<br>(N=41) |  | High ESS<br>(N=31) |  | p-value |
| --- | --- | --- | --- | --- | --- | --- | --- |
|  | N |  | N |  | N |  |  |
| Split |  | 53 (73.6) |  | 30 (73.2) |  | 23 (74.2) |  |
| Body Mass index (BMI), kg/m<sup>2</sup> | 72 | 37.8 ± 9.6 | 41 | 36.5 ± 9.4 | 31 | 39.5 ± 9.9 | 0.19<sup>a1</sup> |
| Epworth Sleepiness Scale | 72 | 2.0 [1.00, 21.0] | 41 | 2.0 [1.00, 2.0] | 31 | 21.0 [20.0, 22.0] | <0.001<sup>b</sup> |
| Total Sleep Time, min | 72 | 319.5 [278.0, 371.0] | 41 | 311.0 [242.0, 359.0] | 31 | 330.0 [304.0, 385.0] | 0.031<sup>b</sup> |
| Apnea Hypopnea Index | 72 | 13.4 [6.0, 38.7] | 41 | 13.1 [6.4, 36.6] | 31 | 13.4 [5.9, 42.1] | 0.66<sup>b</sup> |
| Obstructive Sleep Apnea (Apnea Hypopnea Index, AHI ≥5) | 72 | 61 (84.7) | 41 | 34 (82.9) | 31 | 27 (87.1) | 0.75<sup>d</sup> |
| % Sleep Time with SaO<sub>2</sub><90% | 72 | 1.6 [0.15, 10.6] | 41 | 2.0 [0.30, 8.0] | 31 | 1.00 [0.10, 19.7] | 0.90<sup>b</sup> |
| Arousal Index | 71 | 22.5 [13.6, 39.9] | 40 | 24.9 [15.6, 39.1] | 31 | 19.9 [11.7, 39.9] | 0.44<sup>b</sup> |
| Mean oxygen saturation | 72 | 93.0 [92.0, 95.0] | 41 | 93.0 [92.0, 94.0] | 31 | 93.0 [91.0, 95.0] | 0.58<sup>b</sup> |
| Minimum oxygen saturation | 72 | 85.5 [75.0, 89.0] | 41 | 86.0 [74.0, 89.0] | 31 | 85.0 [77.0, 89.0] | 0.72<sup>b</sup> |
| Sleep Stage % N1 | 69 | 6.1 [3.5, 11.0] | 39 | 6.8 [3.9, 13.4] | 30 | 4.5 [2.6, 7.0] | 0.023<sup>b</sup> |
| Sleep Stage % N2 | 69 | 67.0 [55.3, 74.7] | 39 | 67.2 [55.2, 74.7] | 30 | 66.7 [55.3, 74.7] | 0.96<sup>b</sup> |
| Sleep Stage % N3 | 68 | 5.3 [0.00, 15.1] | 39 | 2.4 [0.00, 16.3] | 29 | 8.4 [0.00, 13.9] | 0.51<sup>b</sup> |
| Sleep Stage % REM | 70 | 17.9 [12.6, 23.4] | 40 | 17.4 [12.5, 20.1] | 30 | 20.1 [13.1, 26.3] | 0.17<sup>b</sup> |
| COVID registry: Coronary Artery Disease | 72 | 8 (11.1) | 41 | 4 (9.8) | 31 | 4 (12.9) | 0.72<sup>d</sup> |
| Hypertension | 72 | 44 (61.1) | 41 | 28 (68.3) | 31 | 16 (51.6) | 0.15<sup>c</sup> |
| Heart failure | 72 | 10 (13.9) | 41 | 6 (14.6) | 31 | 4 (12.9) | 0.99<sup>d</sup> |
| Asthma | 72 | 22 (30.6) | 41 | 11 (26.8) | 31 | 11 (35.5) | 0.43<sup>c</sup> |
| COPD/emphysema | 72 | 13 (18.1) | 41 | 4 (9.8) | 31 | 9 (29.0) | 0.035<sup>c</sup> |
| Cancer | 72 | 11 (15.3) | 41 | 7 (17.1) | 31 | 4 (12.9) | 0.75<sup>d</sup> |
| Diabetes | 72 | 26 (36.1) | 41 | 14 (34.1) | 31 | 12 (38.7) | 0.69<sup>c</sup> |
| Smoking | 72 |  | 41 |  | 31 |  | 0.38<sup>d</sup> |
| No |  | 46 (63.9) |  | 23 (56.1) |  | 23 (74.2) |  |

**Table 1 - Patient Characteristics**
|  | Overall<br>(N=72) |  | Low ESS<br>(N=41) |  | High ESS<br>(N=31) |  | p-value |
| --- | --- | --- | --- | --- | --- | --- | --- |
|  | N |  | N |  | N |  |  |
| Current Smoker |  | 7 (9.7) |  | 5 (12.2) |  | 2 (6.5) |  |
| Former Smoker |  | 19 (26.4) |  | 13 (31.7) |  | 6 (19.4) |  |
| Smoking pack years | 72 |  | 41 |  | 31 |  | 0.092<sup>b</sup> |
| 0.Never smoked |  | 59 (81.9) |  | 31 (75.6) |  | 28 (90.3) |  |
| 0-10 |  | 4 (5.6) |  | 2 (4.9) |  | 2 (6.5) |  |
| 10-30 |  | 8 (11.1) |  | 7 (17.1) |  | 1 (3.2) |  |
| 30+ |  | 1 (1.4) |  | 1 (2.4) |  | 0 (0.00) |  |

Statistics presented as Mean ± SD, Median [P25, P75], N (column %).
p-values: a1=t-test, a2=Satterthwaite t-test, b=Wilcoxon Rank Sum test, c=Pearson's chi-square test, d=Fisher's Exact test.

### Sleep Study Data Acquisition

These studies were conducted in accordance with the American Academy of Sleep Medicine guidelines using Polysmith software version 10 (Nihon Kodhen). For polysomnography (PSG) and split studies, the following signals were recorded: a standard 10–20 EEG montage, electrocardiogram, electromyography of the chin and bilateral anterior tibialis muscle, electrooculography, and pulse oximetry. Oral and nasal airflow were measured with a thermistor and nasal cannula, respectively. Respiratory effort was measured with plethysmography bands at the chest and abdomen, including the summation channel. Sleep staging and respiratory events scoring were conducted based on the American Academy of Sleep Medicine scoring guidelines [9].

### EEG preprocessing

Pre-processing of EEG data, feature extraction, statistics, and machine learning were performed using MATLAB (MathWorks). Since sleepiness is a feature of wakefulness, only EEG during resting state wakefulness were analyzed, defined as the EEG recording time when subjects were awake prior to the sleep study. EEG was collected at a sampling rate of 200 Hz. A high-pass filter with a passband frequency of 1 Hz and a notch filter with a stop-band of 57.5–62.5 Hz were applied to all recordings. All EEG recordings were first visually inspected to confirm signal quality for each channel; channels considered to be of low or irretrievable quality were excluded from the study. Waveforms in each channel were further divided into 1-second epochs, and each epoch was tested for the presence of artifacts using a previously validated method using a support vector machine (SVM) [8]. Epochs containing artifacts were excluded.

### Statistical analyses

We used paired two-tailed t-tests to compare the band-wise PSDs between the High ESS and Low ESS groups [10] (Figure 1 - bottom). We used two-tailed Wilcoxon rank-sum tests to compare the band-wise PAC from the two groups for each of 4 band pairs and 6 channels. We chose a non-parametric statistical test for PAC because values are constrained between 0 and 1 and are therefore less likely to follow a normal distribution, as required by Student’s t-test. Statistical significance was established throughout at p < 0.05. As individual testing was conducted, and the purpose for the statistical testing was primarily the selection of features for subsequent ML (rather than testing of a null hypothesis), there was no adjustment for multiple comparisons [11, 12]. Unlike simultaneous family testing with a joint null hypothesis comprising two or more null hypotheses, individual testing is utilized to make a decision about one null hypothesis. As each test provides only one opportunity to make a Type I error, the alpha level does not require lowering.

**Figure 1:**
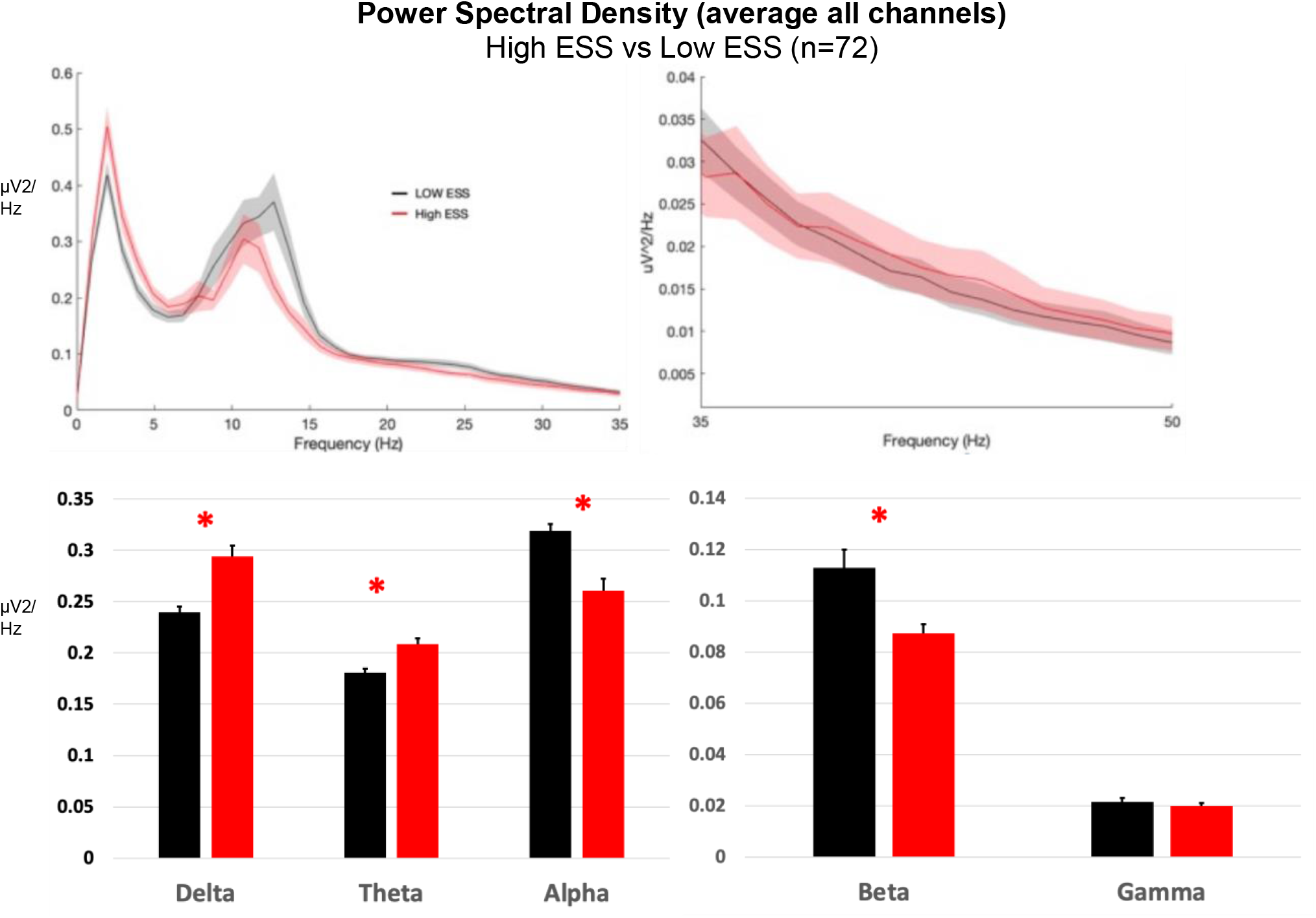
(A) Power spectral density (mean of 6 EEG channels) in High ESS (n=31) and Low ESS (n=41) subjects in the 0-50 Hz frequency range. (B) Mean power in the frequency bands delta (1-4 Hz), theta (5-9 Hz), alpha (10-13 Hz), beta (14-32 Hz) and low gamma (33-52 Hz).

## Results

Our study population was overall middle-aged (mean age 54 years) with a relatively even distribution of men and women and race-based diversity (34.7% African American) and a mild degree of sleep-disordered breathing (apnea-hypopnea index=13.4) Those with a higher degree of EDS were more likely to be slightly younger, female and more obese with longer total sleep time and a lower percentage of N1 sleep stage. Of note, as reflective of our pre-specified design, those with a high degree of EDS had a mean ESS of 21 ±1, in contrast, those without EDS had an ESS=2 ±1.

From the remaining artifact-free epochs of each recording, the following features were extracted: band-wise Power Spectral Density (PSD) for all channels and band-wise Phase-Amplitude Coupling (PAC). To create the band-wise PSD, a periodogram was gathered from artifact-free epochs for each channel, and then these periodograms were averaged together for each channel within each subject. These averaged periodograms were normalized by dividing each frequency bin by the sum of all bins from 3 to 30 Hz. The normalized PSD was used to calculate the band-wise PSD by taking the average of all bins within each of the following four frequency bands: Delta (1-4 Hz), Theta (5–9 Hz), Alpha (10–13 Hz), Beta (14– 32 Hz), and Low Gamma (33–52 Hz). This yielded 5 PSD features for every channel included. The Band by Channel PSD, is similar to the band-Wise PSD. The only difference is that channels are not averaged; each channel is considered separately. This yielded a total of 30 features (5 bands x 6 channels) to be considered. PSD by Hz is similar to Band by Channel, except that instead of bands, we are using bins from 1 to 16 Hz. This yielded to 96 features (16 Hz x 6 channels) to be considered.

PAC was calculated using the Modulation Index (MI) method [13]. The center frequencies used for the phase included all the even numbers from 2 to 20. The center frequencies used for amplitude included all multiples of 3 from 30 to 54. MI was measured for each pair of phase and amplitude frequencies (90 total pairs) for each channel, including only those time points for which there were 5 or more consecutive artifact-free eyes-open epochs. This yielded a 9 × 10 MI matrix, for every channel of every subject, with each row corresponding to one phase center frequency and each column corresponding to one amplitude center frequency. This MI matrix was converted to band-wise PAC for the following 4 pairs of bands: Delta—Low gamma, Theta—Low Gamma, Alpha—Low Gamma, and Beta—Low Gamma. Other band-wise PAC were computed for the following 4 pairs of bands: Delta—Medium Gamma, Theta—Medium Gamma, Alpha—Medium Gamma, and Beta—Medium Gamma. This conversion was accomplished by averaging across the appropriate regions of the MI matrix. This yielded 4 PAC features for each channel and a maximum of 24 PAC features per subject (4 band-pairs x 6 channels).

Bandwise coherence was calculated from each of 15 unique channel pairs by averaging the coherence (MATLAB function mscohere) from each artifact-free epoch during the recordings for both channels in a given pair. This average coherence was divided into five bands in the same manner as PSDs. This yielded 5 coherence features for each channel pair, and a maximum of 75 coherence features per subject (5 bands × 15 channel pairs), coherence values were not computed from channel pairs for which one or both of the channels included artifacts.

Out of the above listed features, 3 features were used to create a feature-set for training binary classification algorithms shown to be significantly different between groups; these included 2 from bandwise coherence, channel pairs O2 – C4 for both Delta and Theta bands, and channel F4 for the 5Hz bin in PSD.

Following recent trends in best machine learning practices [14], several traditional machine learning classification algorithms were considered, using cross-validation and a grid-search strategy to find their optimal hyperparameters. The best results were obtained from k-nearest neighbors algorithm (KNN). To ensure data splits were properly distributed between folds, we validated the classifier using a stratified K cross-validation using *k* = 5. Within our dataset with n=31 for the high ESS group and n=41 for the low ESS group, we computed the following metrics for validation: accuracy, area under the receiver operating characteristic curve (AUC-ROC), precision and recall. Accuracy was calculated within the k-folds cross validation by counting the number of out-of-sample predicted labels that matched the true label of the sample, and dividing this total by the number of samples.

The spectral density for delta, theta, alpha, and beta power bands showed a statistically significant difference in the mean EEG power in High ESS compared to the Low ESS group (Fig 1), including an increase in delta (0.294±0.010 in High ESS, 0.239±0.005 in Low ESS, p<0.001), in theta (0.208±0.005 in High ESS, 0.181±0.004 in Low ESS, p<0.001) and a decrease in alpha (0.260±0.011 in High ESS, 0.318±0.007 in Low ESS, p<0.001) and beta (0.087±0.003 in High ESS, 0.112±0.007 in Low ESS, p<0.001). Analysis of power in individual 6 channels further showed that significant changes were not localized to particular brain areas, except for a distinct increase in alpha in channel F4 (Fig 2). Significant differences in delta, alpha, and beta power bands in each EEG channel were found between low and high ESS groups (Fig 2).

**Figure 2:**
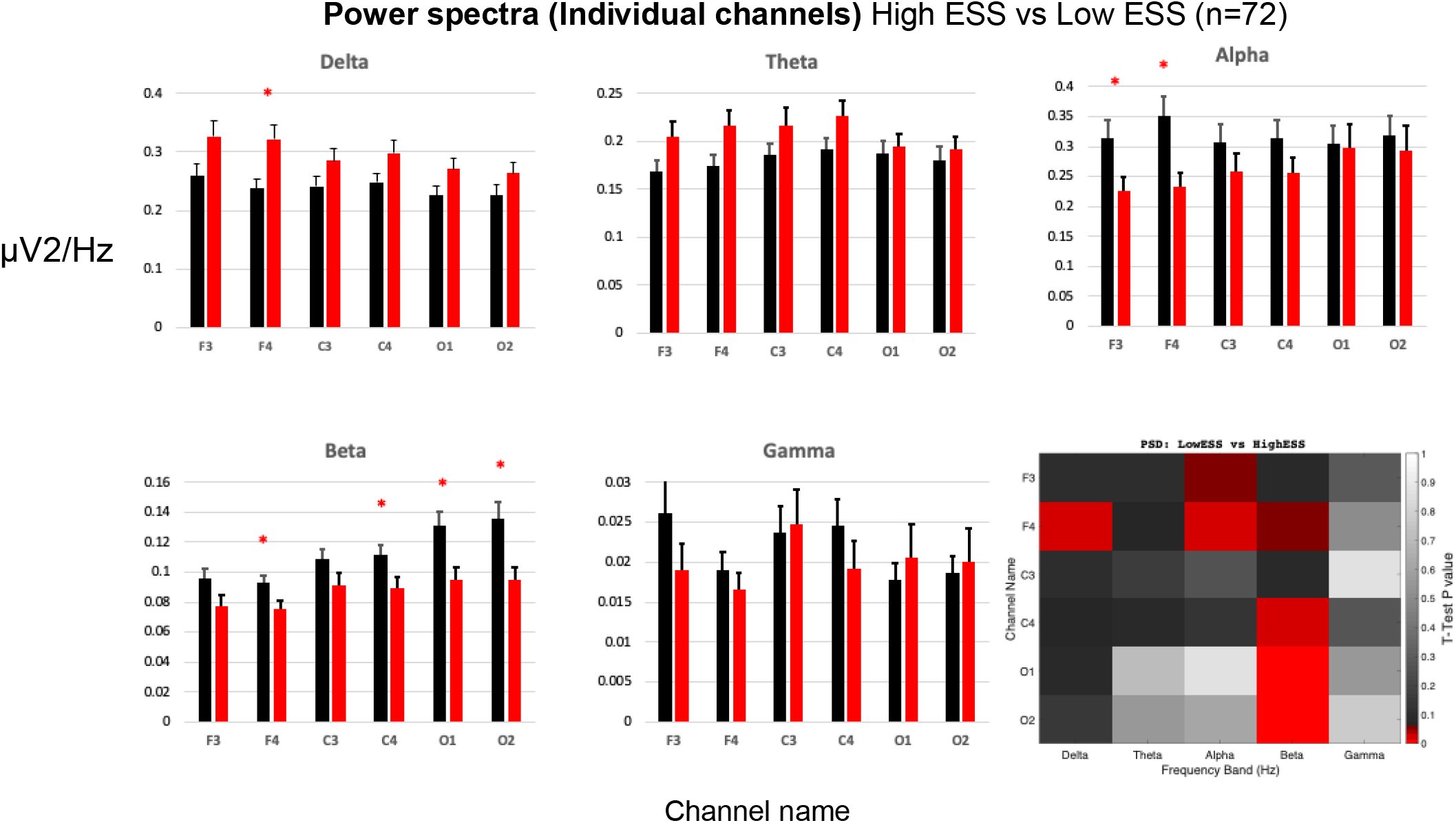
Power spectral density in 6 individual EEG channels in High ESS (n=31) and Low ESS (n=41) subjects in the frequency bands delta (1-4 Hz), theta (5-9 Hz), alpha (10-13 Hz), beta (14-32 Hz) and low gamma (33-52 Hz). Heat map (bottom right) shows t-test values for individual channels in each band (red hue indicates p<0.05).

For Phase-amplitude coupling (PAC) in 6 individual EEG channels, there was no significant difference between low gamma and delta, theta, alpha, beta respectively, as well as between medium gamma and delta, theta, alpha, beta respectively (Fig 3).

**Figure 3:**
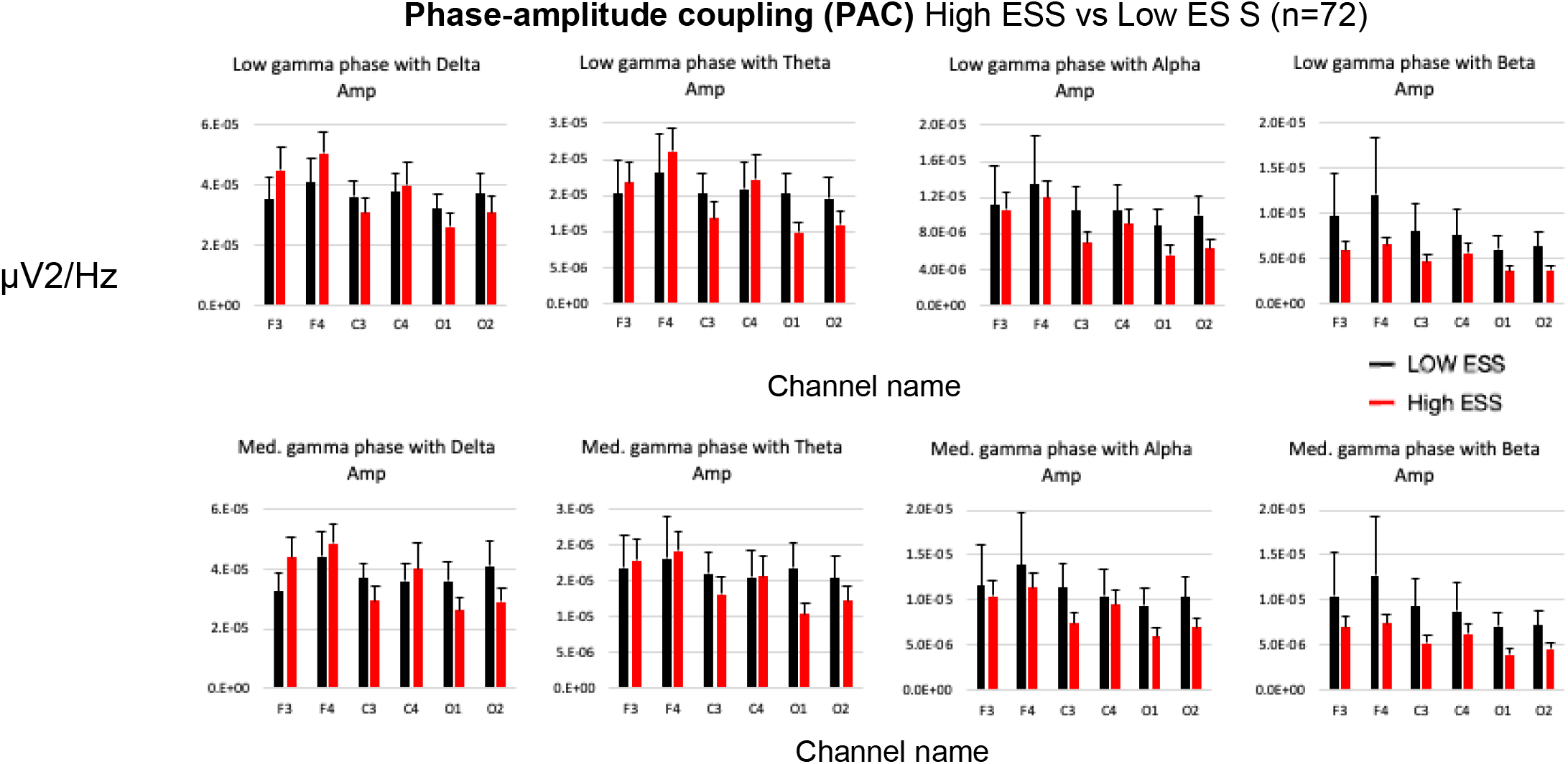
Phase-amplitude coupling (PAC) in 6 individual EEG channels in High ESS (n=31) and Low ESS (n=41) subjects between low gamma and delta, theta, alpha, beta respectively (upper row), as well as between medium gamma and delta, theta, alpha, beta respectively (lower row). No statistically significant difference was noted between groups in any individual channel.

In Figure 4, we compared the coherence in 6 individual EEG channels across each frequency band, significant difference was for the high ESS and low ESS groups and found between the coherence in some of the individual channels like C4-M1 and O2-M1.

**Figure 4:**
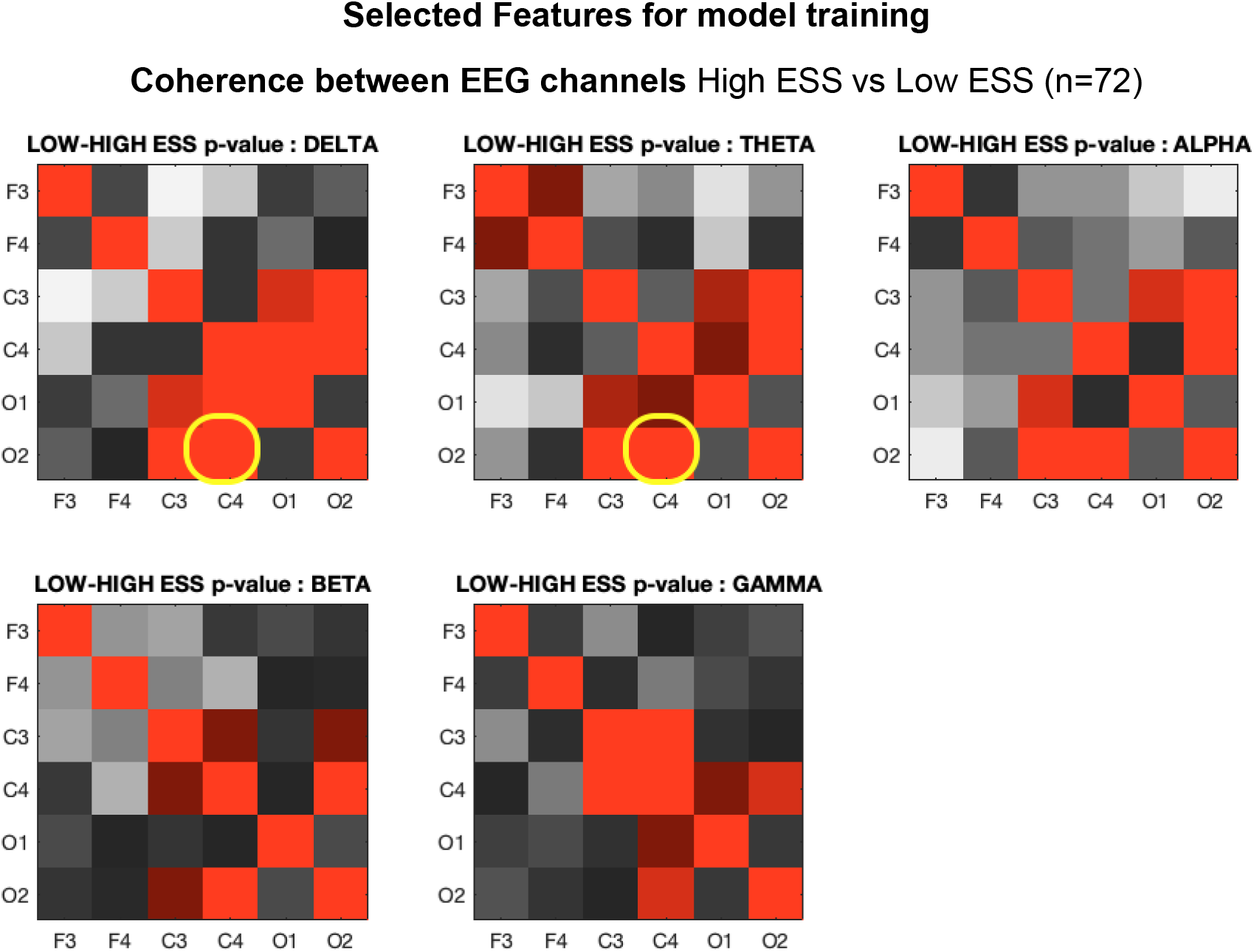
Coherence in 6 individual EEG channels in High ESS (n=31) and Low ESS (n=41) subjects. The values are symmetrical across one diagonal (the 2 highlighted cells were selected for ML).

**Figure 5:**
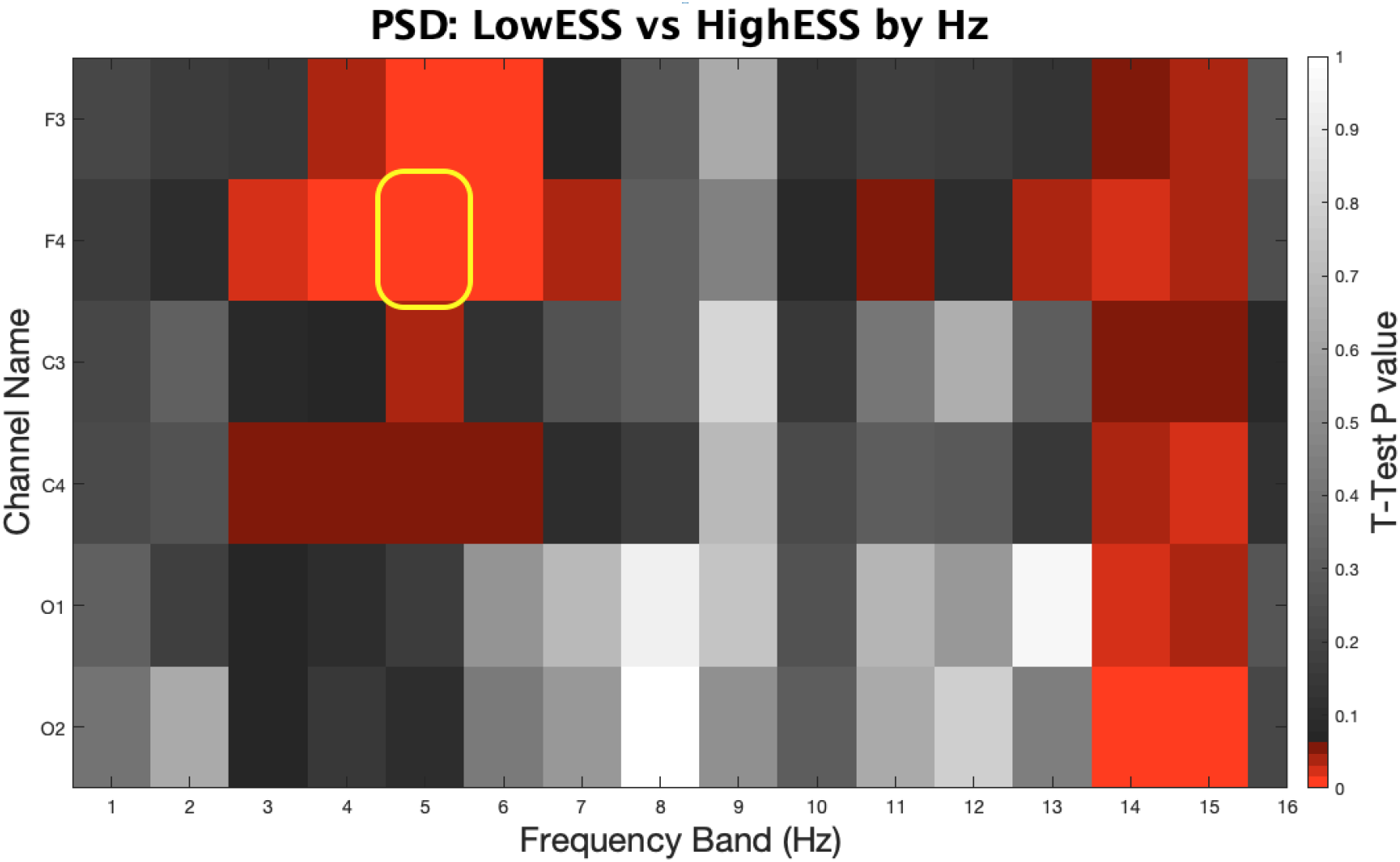
Power spectral density in 6 individual EEG channels in High ESS (n=31) and Low ESS (n=41) subjects in the frequency from 1 Hz to 16 Hz. Heat map shows t-test values for individual channels in each Hz bin (red hue indicates p<0.05) (the highlighted cell was selected for ML).

The relevant features generated from the PSD and Coherence analysis were used to train k-NN binary classifier of high versus low ESS. Our model reached an accuracy of 81.9%, an AUC-ROC of 74.2%, a precision of 84.3% and a recall of 71.0%.

## Discussion

Clinically, EDS overlaps with other common sleep and mental disorders that are considered causal to EDS, such as obstructive sleep apnea (OSA), narcolepsy, depression, and COVID-19 [15], with an estimated prevalence of 20% of adults in the United States [16]. These causes are usually misdiagnosed or undiagnosed in the population, even when a patient performs a sleep study ([17], [7]).

Several approaches have been investigated to develop an objective neurophysiologic biomarker capable of capturing symptoms of EDS. For example, the least absolute shrinkage and selection operator (LASSO) was used to predict ESS from EEG signals collected from train drivers, but with varying degrees of success and requiring more complex computational techniques compared to our study, which was guided by a statistical approach for the selection of MK features [19]. Another proposed sleepiness biomarker is the odds ratio product, computed from the delta, theta, alpha-sigma, and beta frequency bands from EEG signals, and its association to ESS [20]. Despite previous work, however, the direct classification of high or low ESS given an individual EEG signal lacks appropriate rigor and transparency in the ML approach.

In this study, we demonstrate the feasibility of a fully automated analytical pipeline, using resting state EEG during wakefulness and ML, of accurately classifying EDS as low versus high. Our results showed that the average power spectrum across EEG channels in low ESS patients is significantly enhanced in the alpha and beta bands and attenuated in the delta and theta bands compared to high ESS patients.

After analyzing sleep studies in two groups of patients, we identified significant differences in the delta, theta, alpha, and beta frequency bands between those who reported high versus low sleepiness. Aiming to estimate daytime sleepiness at the individual level, we trained a k-NN classifier reaching 81.9% accuracy, 84% precision and 71% of recall. Thus, our work demonstrates a potential for EEG analysis to generate biomarkers for excessive daytime sleepiness, and the use of signal processing and machine learning to classify sleepiness at the individual level.

These results suggest that EEG contains information that could be leveraged to quantitively and reliably assess EDS, thereby enhancing clinical care. We were mostly motivated by the AASM guidelines [18] for sleep staging, in which EEG patterns are highlighted as markers for defining the awareness of an individual, albeit qualitatively.

Some limitations of our work deserve attention. Since the sample size could be considered relatively small (n=72) in the field of EEG/ML [14], our model would benefit from validation against a prospective dataset as per machine learning best practices, as well as across different geographical sites. We also used signal processing techniques to extract EEG features, for example power in predefined delta, theta, alpha, beta, and gamma bands. This may have limited our feature space for biomarkers in contrast to other techniques that based on training deep learning models directly from EEG raw signals [21], although such techniques are non-transparent and more computationally demanding. We also acknowledge that some demographic variables might have an impact on EEG, such as age [22-25] and gender [26, 27], and we did not specifically control for such parameters, although mean demographic values were overall comparable.

Finally, we envision that our fully-automated and quantitative method for assessing EDS can be operationalized by adding our method’s output into patients’ Electronic Health Record after performing a polysomnogram test. We foresee an intervention, such as cognitive-behavioral therapy and sleep hygiene, but more research is needed for proper clinical implementation.

## Conclusion

In this study, we investigate the potential of using EEG signals recorded during the awake period in polysomnograms as biomarkers for excessive daytime sleepiness. We identified significant differences in the delta, theta, alpha, and beta frequency bands between those who reported high versus low sleepiness measured by the widely used ESS. Aiming to estimate daytime sleepiness at the individual level, we trained a k-NN machine learning classifier that reached 81.9% accuracy, 84% precision, and 71% recall in a retrospective cohort. Thus, this study innovates in building a direct association between EEG and ESS. Many areas can be improved, including the problem of underdiagnosed EDS.

